# Human Chr18: “Stakhanovite” Genes, Missing and uPE1 Proteins in Liver Tissue and HepG2 Cells

**DOI:** 10.1101/2020.11.04.358739

**Authors:** George S. Krasnov, Sergey P. Radko, Konstantin G. Ptitsyn, Valeriya V. Shapovalova, Olga S. Timoshenko, Svetlana A. Khmeleva, Leonid K. Kurbatov, Yana Y. Kiseleva, Ekaterina V. Ilgisonis, Mikhail A. Pyatnitskiy, Ekaterina V. Poverennaya, Olga I. Kiseleva, Igor V. Vakhrushev, Anastasia V. Tsvetkova, Ivan V. Buromski, Sergey S. Markin, Victor G. Zgoda, Alexander I. Archakov, Andrey V. Lisitsa, Elena A. Ponomarenko

## Abstract

Missing (MP) and functionally uncharacterized proteins (uPE1) comprise less than 5% of the total number of human Chr18 genes. Within half a year, since the January 2020 version of NextProt, the number of entries in the MP+uPE1 datasets has changed, mainly due to the achievements of antibody-based proteomics. Assuming that the proteome is closely related to the transcriptome scaffold, quantitative PCR, Illumina HiSeq, and Oxford Nanopore Technology were applied to characterize the liver samples of three male donors compared with the HepG2 cell line. The data mining of Expression Atlas (EMBL-EBI) and the profiling of our biospecimens using orthogonal methods of transcriptome analysis have shown that in HepG2 cells and the liver, the genes encoding functionally uncharacterized proteins (uPE1) are expressed as low as for the missing proteins (less than 1 copy per cell), except for selected cases of HSBP1L1, TMEM241, C18orf21, and KLHL14. The initial expectation that uPE1 genes might be expressed at higher levels than MP genes, was compromised by severe discrepancies in our semi-quantitative gene expression data and in public databanks. Such discrepancy forced us to revisit the transcriptome of Chr18, the target of Russian C-HPP Consortia. Tanglegram of highly expressed genes and further correlation analysis have shown the severe dependencies on the mRNA extraction method and analytical platform.

Targeted gene expression analysis by quantitative PCR (qPCR) and high-throughput transcriptome profiling (Illumina HiSeq and ONT MinION) for the same set of samples from normal liver tissue and HepG2 cells revealed the detectable expression of 250+ (92%) protein-coding genes of Chr18 (at least one method). The expression of slightly more than 50% protein-coding genes was detected simultaneously by all three methods. Correlation analysis of the gene expression profiles showed that the grouping of the datasets depended almost equally on both the type of biological material and the experimental method, particularly cDNA/mRNA isolation and library preparation. The dependence on the choice of bioinformatics analysis pipeline was also noticeable but significantly less. Furthermore, the combination of Illumina HiSeq and ONT MinION sequencing to validate proteotypic peptides of missing and uPE1 proteins was performed for the heat-shock factor binding protein HSBP1L1 (missing protein, recently transferred to PE1 category) and uncharacterized protein C18orf21 (uPE1). We observed that a nonsynonymous SNP led to the loss of the site of trypsinolysis in HSBP1L1. The modified version of HSBP1L1 was included in the sequence database and searched against the MS/MS dataset from Kulak, Geyer & Mann (2017), but delivered no significant identification. Thus, HSBP1L1 is still missing for the MS-pillar of C-HPP, although its existence at the protein level has been confirmed.

## Introduction

The Chromosome-Centric Human Proteome Project (C-HPP ^1^) is currently in a mature state with a rough benchmark of 85% of the human proteome covered throughout different tissue/cell types^2^. The remaining portions of the uncovered proteome are challenging because of two issues. First, there are missing proteins never detected by mass spectrometry (MS) or antibodies (Ab)-based experiments at the trustable level of confidence^3^. Second, some proteins were sometimes detected in a certain type of biospecimen carrying no clear function. Although the segregation of the unexplained proteins into two groups is obviously convenient, missing and uPE1 proteins from tissues or cells appear similar due to the equally low level of mRNA expression. Moreover, according to NextProt releases, missing proteins sometimes pass into the uPE1 category, and such transfer confers the ultimate task of the current phase of C-HPP. We hypothesized, that missing and uPE1 proteins might not be so different from the viewpoint of C-HPP mass-spectrometry pillar, particularly if specific biospecimens are analyzed.

For the human chromosome 18 (Chr18, the Russian C-HPP consortium^4^), the baseline metrics to the beginning of 2020 were 265 protein-coding genes (PE1-PE4), among which protein evidence of the level PE1 was available for 252, 13 proteins were missing and 10 had a uPE1 status. This list of 23 proteins (10 PE2, 12 PE3 and just one PE4) became our targets for the neXt-MP50 and CP50 challenges^2^. To capture highlighted targets in this article we relied upon the new sequencing technology of the Oxford Nanopore enhanced by further targeted data mining for the missing and uPE1 proteins.

The absence of some proteins could be reasonably attributed to alterations in the primary structure^5^, when proteotypic peptides can accidentally fall into the splice junctions, or carry nonsynonymous polymorphisms, affecting the peptide retention time and mass-to-charge ratio in LC–MS/MS analysis. Thus, the increased accuracy of transcriptome data became ultimately essential, despite many studies already published on the chromosome-centric transcriptome to proteome mapping, including those from our group^6–8^, from the Spanish Proteome society^9^, reports on Chr9^10^ and Chr17^11^, and also the report on the transcriptome-to-translatome-to-proteome contribution from the Chinese consortium^12^. Inspired by C-HPP work tasks a corpus of bioinformatics tools^13,14^ and databases^15,16^ was developed to manage the transcriptomes. However, little attention was given to the question of the reliability of the underlying transcriptome data. In most cases, the transcriptome was analyzed by a single method—*e.g.,* RNA-Seq—and at best validated by a second method—*e.g.*, by quantitative PCR. For example, SOLiD technology was applied for the whole-transcriptome analysis of HepG2 cell lines and liver tissues^17^ (these biomaterials were proposed by the Russian C-HPP Roadmap^4^) and validated by only 45 selected mRNAs by droplet digital PCR (ddPCR). Some other data included Illumina GaII_x_ and SOLiD profiling of Chr18^7^ but should be considered as outdated due to the novelties in the next-generation sequencing (NGS) platforms.

Validating the transcriptome versus the proteome by quantitatively correlating gene product abundance^18,19^ was an attractive idea. The Chr18 consortium (Russia) investigated such an approach in detail. In capturing the transcriptome to proteome relationships everyone get used to general arguments referring to NGS platform dependence of experiments, gene-to-gene difference in effectiveness of transcription, the rate of protein synthesis and degradation and so on^20,21^, binding all these claims to the whole genome, not to a single chromosome. Contrastively to the whole genome assumptions, in the report^22^ the chromosome-centric transcriptome to proteome correlations were analyzed. The mRNAs levels were determined independently by quantitative PCR (qPCR) of Chr18 genes, and then supported by the next-generation sequencing on Illumina and SOLiD platforms and further by the shotgun MS data. Actually, studies in the field of Chr18 transcriptome profiling and targeted proteome mapping in liver tissue and HepG2 cells^6^ revealed a poor correlation between transcriptome and proteome data. Radko et al.^8^ investigated to which extent the targeted PCR-based transcriptome mining can contribute to the problem of the missing proteins of human Chr18. A summary of these chromosome-centric efforts revealed the unexpectedly low quantitative correlation, with no satisfactory explanation^5^.

To create at least some ground for the analysis of missing and uPE1 species, we reanalyzed the transcriptome of Chr18 to gain more accurate data and assess the level of errors in such data by comparing the results from different platforms of transcriptome quantitation. We recruited the gold-standard method of quantitative PCR (RT-PCR and ddPCR) together with well established HiSeq/Illumina. In addition to these methods, the recently emerged sequencing method articulated itself as the Oxford Nanopore Technology (ONT) was recruited here for C-HPP by using an ONT MinION sequencer^23^, the low-cost portable sequencing machine. The technology produces lengthy reads up to 10^4^ nucleotides advantageously to the Illumina platform, which could obtain reads of 50- to 300-nucleotides long. However, the disadvantage of ONT is that long reads contain errors at the rate of approximately 3-5 lost or misread sites per 100 sequenced nucleotides. ONT sequencing was characterized in genomics by reading up to 70 thousand nucleotides; at transcript level the read length was naturally limited by the mRNA length and quality. For the first time, this technology was applied to the human transcriptome^23^ to analyze seven human cell lines and a set of tissue samples, including that of the human liver. For the LC2/ad cell line, the ONT performance was compared to a short-read RNA-seq data with a pretty good correlation (*r*_p_=0.88) between the methods Authors^23^ thoroughly compared the expression levels of selected genes and found a significant correlation (*r*_p_=0.82) between ONT and qPCR, suggesting the ONT MinION is suitable for the quantitative assessment of the human transcriptome.

Herein, by analysis of 3 samples of healthy livers and hepatocyte-related HepG2 cells, we pursued a double task. First, we compared the three methods of transcript identification and quantification to understand, taking the Chr18 transcriptome as an example, the extent to which the quantitative profiles are consistent within the same biological sample. This question must be answered before explaining the poor transcriptome-proteome correlation. Second, we probed an approach to select promising targets from the set of genes encoding missing and uPE1 proteins by mapping them on the short (Illumina) and long (ONT) reads to explore the occurrence of mutations and/or splice junctions within those coding transcript sequences, which could affect protein detection.

## Experimental Section

### Human liver samples and HepG2 cells

Samples of human liver were collected at autopsy from 3 male donors (designated further as donors #1, #3, and #5) aged 65, 38, and 54 years with the approval of the N.I. Pirogov Russian State Medical University Ethical Committee (protocol #3; March 15, 2018) with the informed consent from donor’s representatives. The donors were HIV and hepatitis free, and the sections had no histological signs of liver diseases. The postmortem resected samples were immediately placed into RNAlater RNA Stabilization Solution (Thermo Fisher Scientific, USA) and stored at −20°C until further use.

HepG2 cells (ATCC HB-8065, ATCC, USA) were grown to approximately 80% confluence and harvested. The cells were washed 3 times with PBS, counted using an EVE automated cell counter (NanoEntek, South Korea), pelleted by centrifugation, and kept in liquid nitrogen until further use.

### Transcriptome profiling using reverse transcription qPCR

For transcriptome profiling with qPCR, total RNA was isolated from liver tissue samples and HepG2 cells using the RNeasy Mini Kit (Qiagen, Germany) according to the manufacturer’s protocol. The on-column DNase digestion step was performed using the RNAse-Free DNase Set (Qiagen, Germany). The isolated total RNA was quantified using a Qubit 4 fluorometer and the Qubit RNA HS Assay Kit (Thermo Fisher Scientific, USA), and the RNA quality was assessed using a Bioanalyzer 2100 System (Agilent Technologies, USA). The RIN numbers for all preparations of total RNA were 7.5 or higher. Synthesis of cDNA was carried out using the AffinityScript qPCR cDNA Synthesis Kit and random primers (Agilent Technologies, USA) according to the manufacturer’s recommendations. The cDNA samples were stored at −20°C until further use.

The amount of each transcript encoded on Chr18 was assessed by measuring the number of copies of pertinent cDNA in the cDNA preparation derived from total RNA. qPCR was conducted in two formats—droplet digital PCR (ddPCR; 49 transcripts) and PCR in real time (PCR-rt; 226 transcripts)—employing the earlier designed set of primers and probes ^22,24^, with minor exceptions. While ddPCR was performed as described previously ^22,24^, the transcriptome profiling by PCR-rt was carried out using the Δ*Ct* method ^25^.

To calculate the copy number of a transcript per cell, the copy number per PCR probe was normalized by dividing it by the amount of total RNA in the PCR probe (200 ng). The number of transcripts per nanogram of total RNA was brought to the copy numbers per cell based on the amount of total RNA in hepatocytes and HepG2 cells, reported to equal 40 pg/cell^26^.

### Illumina HiSeq sequencing and bioinformatics analysis

Total RNA was isolated using the Extract RNA kit (Eurogen, Russia). RNA quality was evaluated using the Bioanalyzer 2100 System (Agilent Technologies). The RIN numbers varied from 7.3 to 9.1. Clustering and sequencing were carried out using the Illumina HiSeq 2500 system (2 lanes per 8 samples) according to the manufacturer’s protocols (Denature and Dilute Libraries Guide; Sequencing in Rapid Run Mode). For each replicate, we derived from 32 to 59 million reads.

The derived fastq files were analyzed by FastQC and then were processed by Trimmomatic. The read mapping and expression quantification were carried out employing STAR 2.7 (splice-aware mapping to genome), bowtie2 (mapping to transcripts), RSEM 1.3 (quantifications of the reads), and Salmon (quasi-mapping and quantification) software packages. The genome GRCh38.p12 assembly (Ensembl release 97) was used as a reference. Finally, we compared the results obtained with STAR-RSEM, bowtie2-RSEM and Salmon, calculated the Spearman/Pearson correlation coefficients and created clustering dendrograms. The distance between samples/pipelines (i.e., dissimilarity rate) was evaluated as 1–corr.coeff. To create dendrograms, we used the complete linkage hierarchical clustering method.

The sequencing data obtained in this study is available at NCBI Sequence Read Archive (BioProject ID PRJNA635536).

### MinION sequencing of HepG2 cells and liver transcriptomes

Total RNA was isolated and characterized as for qPCR analysis. The extraction of mRNA from the total RNA preparations was conducted using the Dynabeads™ mRNA Purification Kit (Thermo Fisher Scientific, USA) following the manufacturer’s recommendations. The mRNA preparations were immediately frozen and stored at −80°C until nanopore sequencing.

Nanopore sequencing was carried out using the MinION sequencer (ONT, UK) with FLO-MIN106 flow cells and R9.4 chemistry and the Direct RNA sequencing kit (SQK-RNA002, ONT, UK). The sequencing libraries were prepared strictly following the manufacturer’s protocol with a single exception: 750 ng of mRNA (poly+ RNA) was used as the input in both samples from human liver and HepG2 cell instead of the recommended 500 ng. The SuperScript III Reverse Transcriptase (Thermo Fisher Scientific, USA) was used for reverse transcription and NEBNext, Quick Ligation Module (New England Laboratories, UK) was for end repair and ligation. The Agencourt RNAClean XP magnetic beads (Beckman Coulter, USA) were employed for nucleic acid purification.

The mRNA from HepG2 was sequenced in a 72-h single run. The output was 0.75-Gb sequenced transcripts (0.766 million reads) with a median length of 1.56 kb. The mRNA from the tissue liver of donor #1 was sequenced for 26 h. The flow cell was regenerated using the Flow Cell Wash Kit (ONT, UK), strictly following the manufacturer’s guidance. Next, the newly prepared sequencing library from the liver mRNA of donor #1 was loaded on the flow cell and a 48-h sequencing run was initiated. The overall output was 1.44 million reads with a median length of 1.37 kb.

The fast5 files produced by MinION were uploaded onto the Amazon Web Services ElasticCloud2 and processed using the GPU-powered (Nvidia Tesla V100) virtual instance p3.2xlarge (8×2.7 GHz vCPUs, 1 GPU) by the ONT-provided basecalling software guppy_basecaller ^27^ with parameters “-flowcell FLO-MIN107 -kit SQK-RNA002”. Further pipelines included the quality control by the MinIONqc.R script, followed by mapping the reads onto the gencode.v32.transcriptome using minimap2 v. 2.17^28^. The overall statistics of alignment mapping was produced using the “samtools stats” command, and the quantitative data were further collected by executing the program Salmon. 0.12/1.1.0 with the command line options “quant -p 8 –noErrorModel” ^29^.

### Proteoinformatics

The MS/MS dataset from the Mann’s group was downloaded from the PRIDE databank (project accession number PXD005141) and searched by the engines embedded into the SearchGUI/PeptideShaker platform^30^ and by MaxQuant^31^ using the same parameters, as previously specified^32^.

The peak lists obtained from MS/MS spectra were identified using OMSSA version 2.1.9^33^, X!Tandem Vengeance (2015.12.15.2)^34^ and MS-GF+ release (v2018.04.09)^35^. The search was conducted using SearchGUI. Protein identification was conducted against a concatenated target/decoy version of the Homo sapiens complement of the UniProtKB (version 2020_04, 42361 targeted sequences). The decoy sequences were created by reversing the target sequences in SearchGUI. The identification settings were as follows: trypsin-specific, with a maximum of 2 missed cleavages, 7.0 ppm as MS1 and 20.0 ppm as MS2 tolerances; fixed modifications: carbamidomethylation of C (+57.021464 Da); variable modifications: oxidation of M (+15.994915 Da), acetylation of protein N-term (+42.010565 Da); fixed modifications during refinement procedure: carbamidomethylation of C (+57.021464 Da); variable modifications during refinement procedure: pyrrolidone from E (−18.010565 Da), pyrrolidone from Q (−17.026549 Da), pyrrolidone from carbamidomethylated C (−17.026549 Da). The peptides and proteins were inferred from the spectrum identification results using PeptideShaker^36^ version 1.16.45. The peptide spectrum matches (PSMs), peptides and proteins were validated at a 1.0% false discovery rate (FDR) estimated using the decoy hit distribution.

The same modifications and m/z tolerance settings (10 ppm for MS1 and 0.01 Da for MS2) were used to perform Mascot (version of the 2010 year) and identyPROT^37^ MS/MS searches.

## Results and Discussion

### 1. Overview of Chr18-coded missing and uPE1 proteins

To characterize the transcription of genes encoding uPE1 and missing proteins (MP), we performed an analysis using the Oxford Nanopore system. In total, 1.5 million reads were obtained for liver cells, and more than 80% of these reads were successfully mapped to the human transcriptome version v32. Data for the same samples were collected using the orthogonal platforms Illumina (hereafter also called HiSeq) and quantitative PCR.

The transcripts of the uPE1 proteins were expected to be detected at more than the transcripts encoding the missing proteins. However, the experimental data indicated the opposite: the level of protein transcription in both groups was comparably low. For example, for the uPE1 proteins TMEM200C and KLHL14, 15 TPM and 7 TPM were observed, respectively. The average expression level (for the values: TPM>0, n=8) was 4.48±2.67 TPM and 4.69±1.17 TPM for transcripts encoding missing and uPE1 proteins, respectively.

We compared the values of the expression level of the studied group of genes (MP+uPE1) with the data obtained using alternative methods (Supporting Information Table S1). We confirmed, using both Illumina HiSeq and quantitative PCR, that the transcripts are expressed at an extremely low level, close to the detection limit. Thus, regarding the transcriptome of the studied biomaterials—that is, the liver and HepG2 cells—the genes encoding missing and uPE1 proteins were similar in the expression level and could be analyzed together.

The use of the EMBL-EBI’s gene ExpressionAtlas^38^ database revealed that the genes encoding the missing proteins are not expressed at a significant level in any of the analyzed tissues. For example, the highest values for the ELO3B gene reached a negligible level of 2 TPM in the choroid plexus, and HMSD, the most expressed gene, reached 11 TPM in the spinal cord and 14 TPM in transformed lymphocytes. However, genes encoding uPE1 proteins in some tissues reached 3–5 times higher values than the maximum values for missing protein transcripts: ANKRD29 reached 71 TPM in the lymph nodes or up to 57 TPM for a gene in the thyroid gland.

Confirmation of the above observations was found in ProteinAtlas. In the liver tissue and HepG2 cells, the level of transcription of the uPE1 coding genes and missing proteins was comparatively low; however, in at least 1–2 tissues, the uPE1 proteins were expressed pronouncedly at the level of several dozen TPMs. An interesting observation was the TMEM241 gene, which, according to the data of RNA-Seq from ProteinAtlas, demonstrated little expression in the HepG2 cell line at the level of 5 TPM, producing zero values in all the samples by both sequencing methods used herein but was detected as low as 2.5 copies of cDNA per cell using ddPCR.

The pronounced differences between uPE1 and missing proteins were observed in the NextProt portal and the associated PeptideAtlas resource: the uPE1s were usually highly populated using unique natural peptides, with little or no missing proteins. Notably, one missing protein (CTAGE1) from the number of missing proteins does not have unique proteotypic peptides during tryptic hydrolysis. Two other proteins demonstrated one proteotypic peptide each: the ANKRD30B protein peptide was found in prostate cancer samples^39^, and the proteotypic peptide for HSBP1L1 was found in HepG2 cell samples^32^.

Comparison of the NextProt releases from January and July 2020 revealed the transfers of proteins encoded by human Chr18. The SMIM21 protein has moved from the missing protein category to the uPE1 category. SMIM21 is characterized as “expression level 3” according to ProteinAtlas data, at a high level of normalized expression of mRNA NX=5.0 in the brain cancer line. Another protein, encoded by the C18orf65 gene, changed its category from PE2 to PE5, leaving the ranks of the missing proteins of chromosome 18.

For the HSBP1L1 protein (74 a.a.r.), PeptideAtlas suggests one proteotypic peptide 24 a.a.r. long. It was detected in HepG2 and LNCaT cell lines by Professor Mann’s group^32^ using the deep proteome coverage (8–12 thousand protein groups in the identification list) with original SPIDER technology of low-loss digest mixture separation. Information has been published about the involvement of this gene in the development head and neck cancer^40^, pancreas adenocarcinomas^41^. HSBP1L1 is involved in chemical hepatocyte injury^42^, prioritizing this target to investigate the genetic aspects of liver diseases. In the July 2020 version of NextProt, this protein was transferred from the PE2 (e.g. missing proteins) to PE1 category, with the functional annotation at the “silver” level. To our knowledge, this transfer is not yet confirmed by mass spectrometric data in accordance with the criteria of version 3.0 of the guidelines^43^, at the same time MS-based support was delivered by top-down proteomics^44^ in HeLa cells.

So, at the time of this writing, the updated version of NextProt included 9 missing genes and 11 genes characterized as uPE1 from Chr18.

A special case is presented by the KLHL14 (uPE1) gene, which, according to different methods, was not expressed in all biosamples, except for the HepG2 sample, where its expression level measured by quantitative PCR was reported as 116 copies per cell. The expression level of the housekeeping gene ATP5F1A was comparable — 324 copies per one HepG2 cell on an average.

To rely on transcriptome data for the analysis of MP and uPE1 proteins, all of which exhibit low transcription levels, it is necessary to confirm the reliability of the basic data. In view of this assumption we considered the datasets in two ways: comparison of the list of highly expressed genes – “Stakhanovite genes”^45^ and deciphering a degree of correlation between analytical methods. “Stakhanovite genes” are named after the Soviet coal miner Alexey Stakhanov renowned for his outstandingly hard everyday work.

### 2. “Stakhanovite genes” of Chr18

Missing and uPE1 proteins encoded by the human Chr18 are characterized by low expression of corresponding genes in the liver and HepG2 samples. To estimate the contribution of error in the sample preparation and measuring methods, we started not from the low expressed transcripts, but, oppositely, from the extremely high abundant ones produced by “Stakhanovite genes” (Fig. 1). The choice of significantly expressed genes is justified, as estimations obtained from highly expressed species are prone to noise, thus providing a minimal level of errors.

**Fig. 1.**
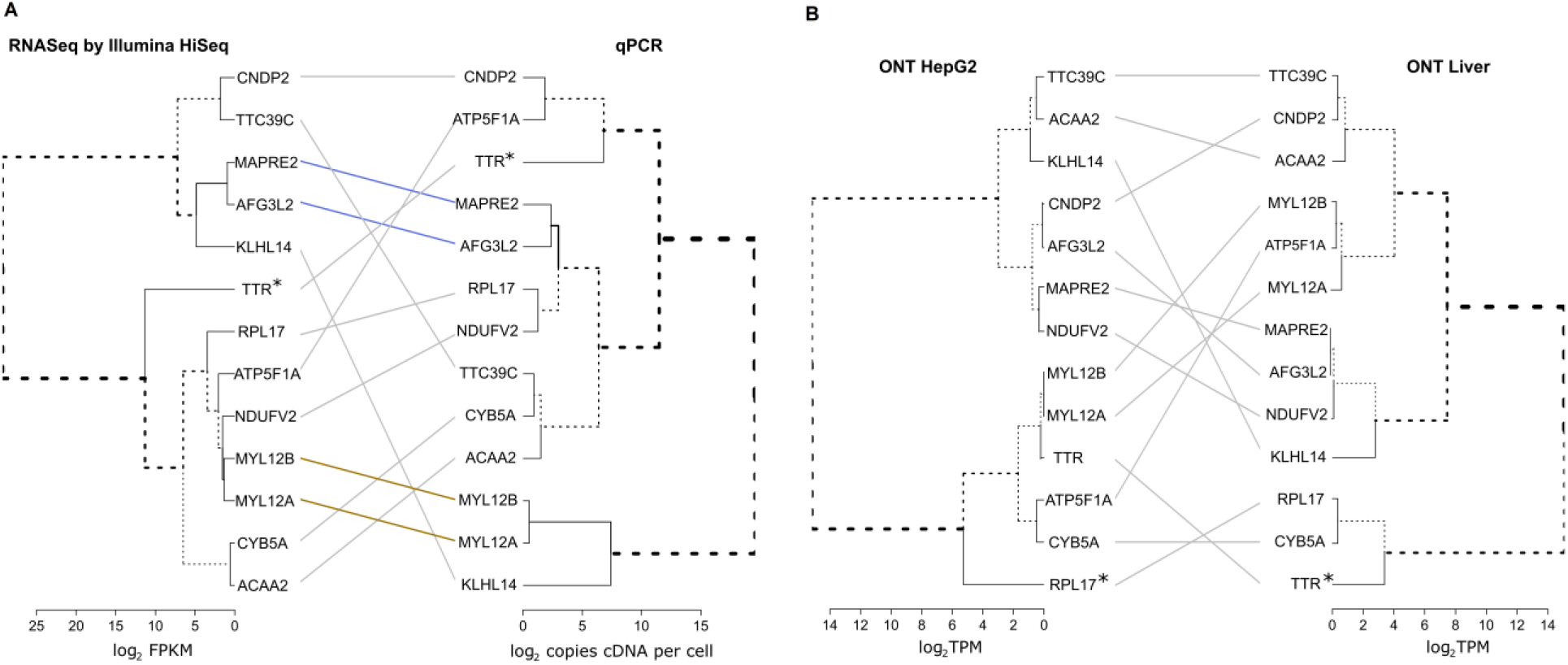
Tanglegram of the most heavily expressed “Stakhanovite” genes with (a) different methods and one sample (human liver) and (b) by the same method, applied for different biospecimens (liver versus HepG2 cell line). Dendrograms were obtained using differences between the estimations of the expression levels as measured by qPCR, Illumina HiSeq and ONT methods and scaled for convenient representation. * indicates the most highly expressed genes. Dendrograms were built using Ward’s clustering and the Euclid distance between log2-transformed data. The Dendextend package^46^ was used to draw tanglegrams and compute entanglement.

Supporting Information Table S2 shows five of the leading “Stakhanovite genes” for each sample studied, exhibiting the highest levels of expression as revealed by ONT MinION, Illumina HiSeq, and qPCR analysis. These 5 topmost genes are responsible for a substantial part of the total mRNA abundance. For example, in the case of ONT MinION sequencing, more than 95% of the total mRNA abundance was attributed to the corresponding “Stakhanovite genes” in both HepG2 cell and liver samples. In this case, the RPL17 gene encoding 60S ribosomal protein L17—a component of the ribosomal complex—leads the list of “Stakhanovite genes” for HepG2 cells. The gene encoding transthyretin, TTR, was expectedly most highly expressed in the liver, which secretes blood plasma ingredients. In addition to RPL17, other housekeeping genes were revealed by ONT sequencing: MYL12A and MYL12B encoding chains of the motor protein myosin; ATP5F1A encoding the ATP-synthase subunit alpha responsible for the energy generation in mitochondria; and CYB5A encoding cytochrome b5, the membrane-bound electron carrier. Among these genes, 4 top genes were common for HepG2 cells and liver tissue, showing a considerable level of qualitative concordance between the biological samples of different origin.

Illumina HiSeq sequencing also revealed some level of qualitative concordance among “Stakhanovite genes” for HepG2 cells and liver tissue samples: TTR and RPL17 were expressed in all liver samples and HepG2 cells, NDUFV2 as well as MYL12A were present in HepG2 and at least in two of three liver samples (Fig. 1 and Supporting Information Table S2). The similar level of qualitative concordance among “Stakhanovite genes” was found between the liver samples studied. For qPCR analysis, a relatively high level of matching was observed for the subsets of “Stakhanovite genes” among liver tissue samples: a coincidence of 4 of 5 or even 5 of 5 among the lists of highly expressed genes. Surprisingly, the level of concordance between the subsets of “Stakhanovites” revealed by two sequencing platforms, ONT MinION and Illumina HiSeq, was rather moderate: only 3 items were shared between HepG2 cells and the liver sample; TTR, RPL17, MYL12A and CYB5A were among them. A similar situation was observed for the liver sample when the ONT MinION or Illumina HiSeq data were compared with the qPCR data: only 3 of 5 top genes coincided. When the qPCR data were compared with the HiSeq dataset for the liver tissue samples of donors #3 and #5, only 2 matching hits were identified (out of 5 selected “Stakhanovite genes”). Moreover, for HepG2 cells, no concordance was found between the subset of “Stakhanovite genes” revealed by qPCR and those revealed by either the ONT MinION or Illumina HiSeq sequencing (Supporting Information Table S2).

The leaves of the dendrograms in Fig. 1 correspond to the highly expressed genes of Chr18 (topmost 13 expressed genes were taken for this analysis, each showing itself at least once in the Supporting Information Table S2). Clustering was performed based on the distance metrics, which correspond to the difference in the values of gene expression; therefore, the closer were leaves at the tree, the more similar were the estimations of expression (either FPKM, copies per cell or TPM).

Fig. 1A represents the results of the analysis using two different methods for one and the same liver sample. The leaves were severely mixed between clusters, which were visually observed as crosses between edges connecting two dendrograms. Among the 10 probed genes, only two pairs preserved their closeness in both dendrograms: MAPRE2 neighboring AFG3L2 and subcluster MYL12 subunits A and B. No such neighboring was observed when comparing different samples by a single method, the ONT (Fig. 1B). The assessment of entanglement as a measure of concordance between dendrograms produced similarly low values of 0.31 and 0.28 for Fig. 1A and 1B, respectively. Thus, working within a single, not-so-large chromosome, even for the case of highly expressed genes, the relationship between the quantitative estimations of the expression level was highly discordant. Although this finding may be expected for the samples of normal liver and HepG2 cells derived from hepatoblastoma, it was rather surprising to find such disparity in the results acquired from the same sample analyzed using different methods.

### 3. Correlations of chromosome-centric transcriptome datasets

Different methods of transcriptome analysis may give different results^7,17,47^. At the genome scale, the frequency of RNA-Seq transcripts provides clues for MS identification of the corresponding protein^48^. However, the feature of chromosome-centric quantitative transcriptomics (presumably of quantitative proteomics as well) is that when number of genes is reduced compared with that in the genome, the admissible correlations can be severely affected. This can be especially true for chromosome 18 with a rather small number of genes.

Regarding qPCR, 235 of 265 transcripts encoded on Chr18 were identified in both HepG2 cells and liver samples of each of three donors, while 16 transcripts were detected neither in HepG2 cells nor in liver samples of any of the donors (Supporting information Table S1 and Fig. S1). Furthermore, 14 transcripts were not found in HepG2 cells but were observed in a liver sample of at least one donor, and 10 transcripts were present in HepG2 cells but not in a liver sample of at least one donor (Supporting information Table S1). For common transcripts of HepG2 cells and the liver samples, a good correlation was observed between the levels of the Chr18 log-transformed gene expression values (Supporting information Fig. S2.1, panels A to C). The values of the Pearson correlation coefficient, *r*_p_, are within the narrow interval of 0.76–0.79, agreeing well with the previously obtained *r*_p_ value of 0.78 for the ‘HepG2 cells *vs.* pooled liver sample’ correlation^22^. At the same time, the correlations between the transcript abundances in the liver tissue between donors were found to be strikingly high: *r*_p_ = 0.963–0.966 (Supporting information Fig. S2.1, panels D to F). Similar to qPCR data, a higher correlation was observed between gene expression profiles derived with Illumina HiSeq (log-transformed FPKM values) in liver tissue samples – *r*_p_ = 0.827-0.925 (Supporting information Fig. S2.2). Liver gene expression, quantified by the Illumina HiSeq, was also well correlated with that in HepG2 cells: *r*_p_ values were within the interval of 0.691–0.812 and agreed with those obtained by qPCR profiling (Supporting information, Fig. S2.2 *vs.* Fig. S2.1).

Compared with the short-read sequencing such as Illumina, the long-read ONT sequencing is a relatively new technology. It should be considered as orthogonal to HiSeq, and different bioinformatics analyses are eventually required to treat the raw data. Although the LAST and BWA algorithms (originally developed for the mapping of short reads^49,50^) were also employed for ONT data processing^23^, the minimap2 tool^51^ is nowadays widely used for long noisy read mapping.

Thus, we prepared three datasets from the orthogonal methods and performed correlation analysis supplied with a cluster dendrogram (Fig. 2, Supporting Information Table S3).

**Fig. 2.**
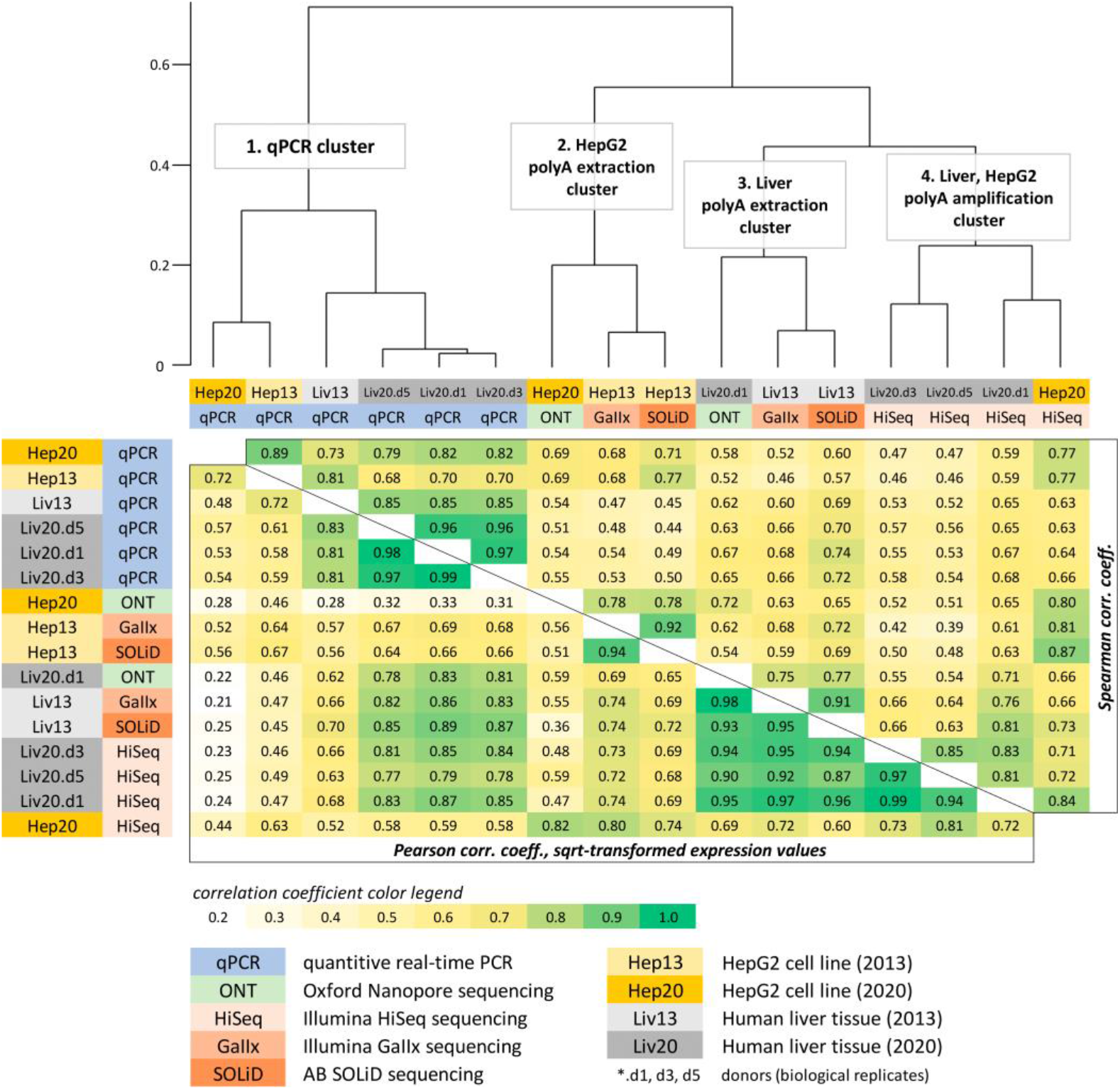
Cross-correlation matrix between gene expression datasets obtained for liver tissue and HepG2 cell line samples from different sources and harvested in our previous work in 2016 (Hep13, Liv13) and current work (Hep20, Liv20). The indices d1, d3 and d5 for Liv20 indicate the particular individuals (donors of postmortem liver samples). SRA datasets: Hep13 - SRX395473 (2013) and SRX390071 (2014); Liv13 - SRX267708 (2013).

The upper triangle of the matrix in Fig. 2 contains the Spearman correlation coefficient between the gene expression values evaluated by qPCR (number of cDNA copies per cell), Illumina HiSeq, and ONT. Additionally, we calculated Pearson correlation coefficients (lower triangle of the matrix; Fig. 2, Supporting information Table S2) for the expression values, which were the square root transformed to reduce the dominant effect of highly expressed genes. The dendrogram was created based on the Spearman rank correlation coefficients (hierarchical clustering using Ward’s D2 method).

In this study, we also included our previously obtained results^24^, which were derived using Illumina GaII_x_ and Applied Biosystems SOLiD platforms for the HepG2 cell line (denoted as Hep13 in Fig. 2; originating from a different source than the HepG2 cells used in the current study) and from the pooled sample of three human livers (denoted as Liv13).

As can be seen from the dendrogram (Fig. 2), the gene expression profiles are clustered into four groups. The division occurs at the level of the methodology: the first cluster was attributed to quantitative PCR method, while other three clusters accommodate the results of next-generation sequencing. Furthermore, to a lesser extent, in the case of qPCR, a separation occurred according to the type of biological material. In the case of sequencing, a separation occurred according to both the biomaterial type and cDNA/mRNA library preparation method. These findings are presented more clearly in the Supporting information Table S3, which includes data on technical replicates and various RNA-Seq data processing pipelines. At the same time, clusters 2 and 3 combined the data from various platforms—Illumina GaII_x_, AB SOLiD, and ONT MinION. These libraries were prepared by polyA mRNA isolation using magnetic microbeads. Finally, cluster 4 included data on HepG2 and liver samples prepared using polyA amplification.

One possible reason for this discrepancy between library prep techniques may be a difference in RNA integrity. Low initial RNA integrity as well as an improper sample preparation procedure induces a shift in the density read coverage towards the 3′-tail of transcripts when using polyA isolation/amplification methods. Thus, we evaluated the differences in the distribution of 5′-to-3′ read coverage density and fortunately found no significant differences between cluster 4 and clusters 2/3 (*i.e.*, between polyA mRNA isolation using the microbead technique and polyA amplification using MINT reverse transcriptase). The data in Supporting information Table S3 provides information on how fragile are dependencies between samples, sample preparation methods, bioanalytical platforms and bioinformatics pipeline of transcriptome data processing. Such dependencies are usually ignored at whole-genome level. The chromosome-centric current phase of discovery of the missing and uPE1 proteins appeals to re-think the paradigm at the transcriptome level.

With several exceptions, the Spearman (rank) and Pearson correlation coefficients were concordant. The highest Spearman’s rank correlation (r = 0.70–0.97) was observed for the qPCR methods (cluster 1). Within this cluster, there were two biomaterials— specific subclusters—within which the data were year dependent as Liv13 segregated from Liv20. In general, the observed correlation pattern reflected that qPCR works much better with low-expression genes than with RNA-Seq mainly because of the stochastic noise and discrete nature of gene expression values derived with RNA-Seq (*i.e.*, read counts).

Notably, different clustering methods produced different results: preferred grouping was observed either by biological material or method. However, the close relationship of the technical replicates and bioinformatics pipelines remained invariant in the dendrograms. Nonetheless, we did not observe a perfect match between the results obtained using various genome annotations (RefSeq/Ensembl/Gencode) and data processing protocols (STAR+RSEM, bowtie2+RSEM, Salmon), as well as between technical replications: the Spearman correlation coefficient between various data processing pipelines or technical replicates ranging from 60% to 97%.

### 4. ONT-oriented selection of missing and uPE1 proteins

We observed a large variability among gene expression values derived by RT– qPCR, Illumina and ONT sequencing from the same biological sample. However, the present work was not intended to clarify the underlying reasons for this variability, but we aimed to highlight that a relatively high correlation between various datasets derived by whole-transcriptome analysis comes primarily from the large dynamic range of transcript abundances. Despite the sampling was confined by a particular chromosome (Chr18, 265 genes), the dynamic range of transcript abundances was still fairly large. Nonetheless, the general correlations may decline (compare the values of correlation coefficients in Fig. 4 with those of 0.92 or 0.9, reported for 18000+ genes by Tyakht *et al.*^17^ or van Deft *et al.*^52^, respectively). For a more limited set of genes—even if these genes are highly expressed such as in the case of “Stakhanovite genes”—obvious disagreement occurred in the levels of their expression measured by the orthogonal methods (tanglegram in Fig. 1A, see also Supporting Information Table S2). The dendrogram topologies (the branching pattern relationship among individual genes) were quite different for the RT–qPCR and Illumina methods (Fig. 1B). For a set of low-abundance transcripts, one should expect a more profound “noise effect”—the apparent well-known fact from whole-transcriptome analysis. Additionally, the number of low-abundance transcripts included in further analysis is heavily affected by TPM and FPKM threshold values set for ONT and Illumina methods, respectively. For example, taking the TPM/FPKM threshold values as ≥ 0.1 resulted in ~60% of Chr18 transcripts detected by all three methods (Supporting Information Fig. S1), while setting the Illumina’s FPKM threshold as > 0.1 would result in less than 50% of such transcripts.

Because we learned from the Expression Atlas (EMBL-EBI) generally and from our own data particularly, that the target missing and uPE1 proteins were present in low copies, we focused ourselves on the presence of a corresponding gene product in all samples, detected by all methods employed. The lower is the level of expression, the more nonrandom would be such an observation. Therefore, in the low-copy range, correlation analysis should be replaced by the unraveling of rarely observed events. This enabled us to perform a simple selection of priorities for the MP/uPE1 array as follows.

From the compendium of the ONT-detectable genes, we selected those that exhibited nonzero values of TPM in both types of biomaterials—*i.e.,* the liver and HepG2. Two missing proteins and four proteins with the uPE1 status were obtained (Table 5). The following information was taken from the neXtProt in Table 5: number of amino-acid residues of the proteoform designated as Iso1 (canonical sequence, often the longest one), number of variant records reported for the given protein as single-amino acid polymorphisms, and number of isoforms (assuming that these could be splice forms or processed forms). Table 1 was completed by indicating for the selected set of genes the FPKM values (Illumina HiSeq), number of cDNA copies (qPCR), and TPM values (ONT MiniION).

**Table 1.** Selected genes of human chromosome 18. First, two genes were annotated as encoding the missing proteins (MPs), while four other genes were attributed to the category of functionally uncharacterized but translated to the protein level with high evidence (uPE1). The neXtProt knowledgebase was used to obtain information about the gene status, number of amino acid residues in the sequence of the isoform 1 (the longest one), single amino acid polymorphisms (SAPs) according to the dbSNP and COSMIC datasets, and the known splice forms (Iso1, Iso2, Iso3).

| Gene Name | HMSD_ | HSBP1L1 | ANKRD29 | C18orf21 | KLHL14 | TMEM200C |
| --- | --- | --- | --- | --- | --- | --- |
| Protein Name | Serpin-like protein HMSD | Heat shock factor-binding protein | Ankyrin repeat domain-containing protein 29 | UPF0711 protein C18orf21 | Kelch-like protein 14 | Trans-membrane protein 200C |
| NextProt AC | NX_A8MTL9 | NX_C9JCN9 | NX_Q8N6D5 | NX_Q32NC0 | NX_Q9P2G3 | NX_A6NKL6 |
| status | MP | MP | uPE1 | uPE1 | uPE1 | uPE1 |
| # of aa in Iso1 | 139 | 75 | 301 | 220 | 628 | 621 |
| variants (SAPs) | 111 | 56 | 231 | 166 | 385 | 648 |
| splice forms | Iso1 | Iso1 | Iso1, Iso3 | Iso1, Iso2 | Iso1, Iso2 | Iso1 |
| qPCR (cDNA copies per cell) |  |  |  |  |  |  |
| Liver #1 | 0.00** | 0.88 | 0.02 | 0.37 | 0.02 | 0.33 |
| Liver #3 | 0.01 | 0.47 | 0.10 | 0.33 | 0.03 | 0.42 |
| Liver #5 | 0.02 | 0.59 | 0.07 | 0.22 | 0.01 | 0.52 |
| HepG2 | ND* | 1.10 | 0.02 | 2.29 | 116.38 | 0.08 |
| Illumina HiSeq (FPKM) |  |  |  |  |  |  |
| Liver #1 | ND* | 3.50 | 0.00* | 4.5 | 0.00** | 0.00** |
| Liver #3 | ND* | 2.50 | 0.50 | 3.0 | 0.00** | 1.00 |
| Liver #5 | ND* | 6.00 | 0.00* | 1.5 | 0.00** | 2.00 |
| HepG2 | ND* | 6.50 | 0.00* | 17.5 | 2.50 | 0.00** |
| ONT MinION (TPM) |  |  |  |  |  |  |
| Liver #1 | 0.21 | 6.31 | 0.13 | 6.20 | 0.00** | 0.04 |
| HepG2 | 0.00** | 6.94 | 0.00** | 16.00 | 1.11 | 0.00** |

\*No reads observed \*\*Rounded to the second decimal place

Table 1 enabled us to select interesting cases of the HSBP1L1 missing protein and C18orf21, the uPE1 protein. Compared with other proteins, these two proteins were covered by ONT reads at modest but detectable levels. By contrast, for example, the serpin-like protein HMSD was poorly represented by the ONT-derived transcripts and undetected by the HiSeq NGS platform. Because of negligible detection, the uPE1 proteins KLHL14 and TMEM200C were discarded. Notably, KLHL14 showed an unexpected finding: the qPCR estimation was 116 copies per cell, including it in a list of “Stakhanovite genes” (Fig. 1).

As of January 2020, the heat shock factor-binding protein (HSBP1L1) was still missing, probably because of its tiny size of only 75 amino acid residues. Additionally, HSBP1L1 was constitutively expressed in the liver and HepG2 cells at the level of approximately 6–7 transcripts per million (roughly corresponding to 1 cDNA copy per cell). The values of FPKM were at the level of 3.5 for liv.d1, and, remarkably, this protein was also detectable in other samples, *viz.* liv.d3 and liv.d5.

The second selected candidate was UPF10711 protein, which is encoded with C18orf21 gene. According to the neXtProt record Q32NC0, this protein was characterized by two isoforms, Iso1 (220 aa) and Iso2 (first 88 aa of Iso1 were missing from Iso2), or three computationally mapped potential isoforms. The ONT approach enabled detection in the liver at the levels of transcription of 0.67 and 4.85 TPM units for the isoforms with accession numbers NX_Q32NC0-1 and NX_Q32NC0-2, respectively.

Illustratively, regarding C18orf21, the Sashimi plot was produced to depict the splicing junctions observed by ONT MiniION and Illumina HiSeq technologies (see Supporting information Fig. S3). Four exons were observed, connected by three solid splice junctions with certain alterations of splice junctions between the first and second exons. The sequence of the proteotypic peptide was observed at the third exon, being safe from the splicing perturbations. We used the standard file of the transcriptome data, which indeed cannot resolve the unexpected splicing events. The deficiency of splicing information should be seriously considered at the current and forthcoming phase of C-HPP.

### 5. Mapping Proteotypic Peptides of HSBP1L1 onto the Transcriptome

In Fig. 3, two proteotypic peptides for the HSBP1L1 protein are shown, the first comprising 24 amino acid residues (Pept-24) and the second 13 a.a.r. (Pept-13). The longer peptide was identified previously^32^ to encounter 20 y and b fragment ions in its spectra obtained in HepG2 cells. When loss-less fractionation was performed in that study for deeper proteome coverage, the deamidated version of Pept-24 was registered in LNCaP cells, with lower coverage by only 10 y- and b-ions. The intensity of the precursor ion in HepG2 was 2 times higher than the signal captured in LNCaP cells. Thus, HSBP1L1 identification by a pair of unique peptides is possible, but not in LNCaP or any of the other 11 cell lines probed earlier^32^

**Fig. 3.**
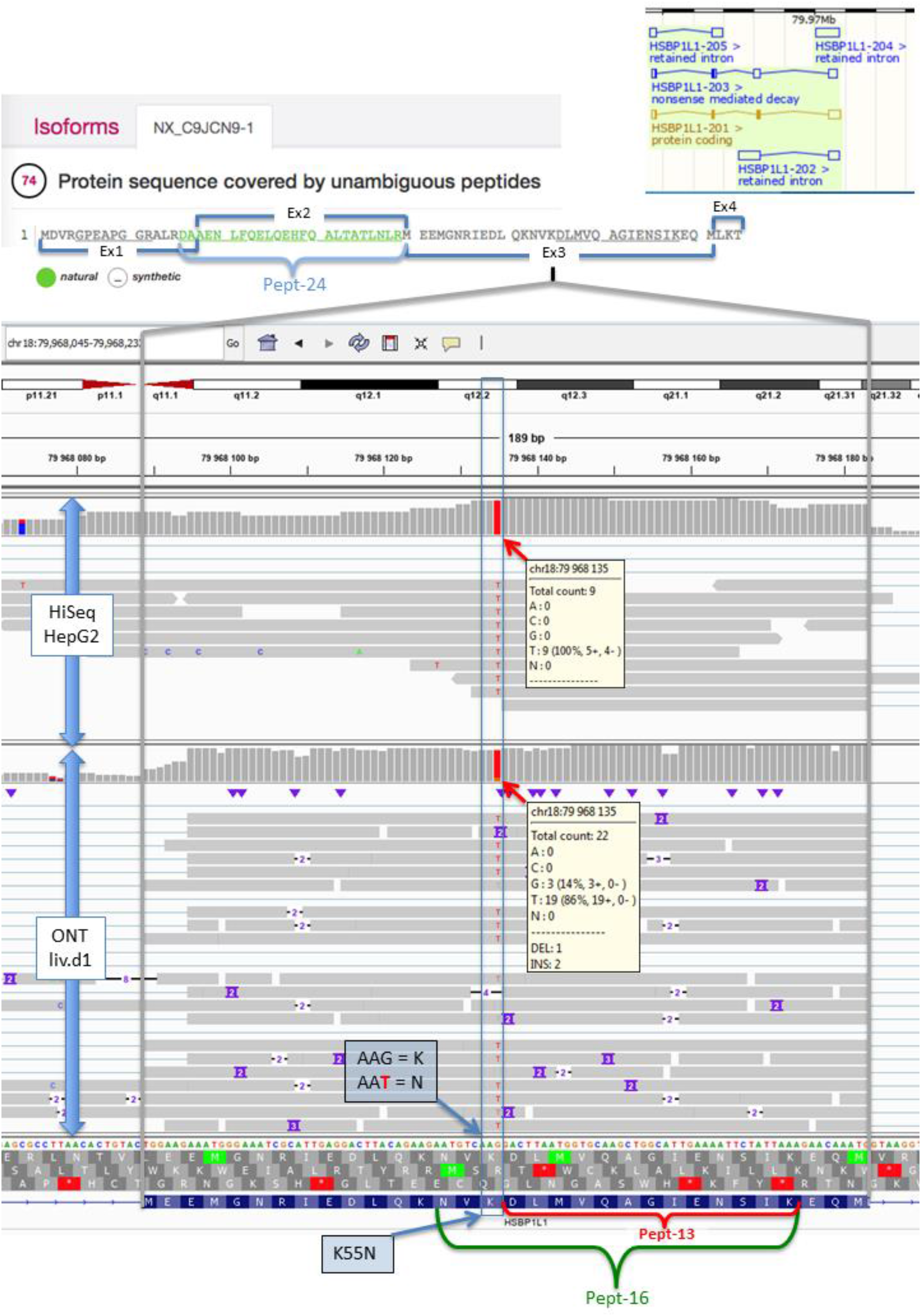
The exon 3 of HSBP1L1 gene (encodes for missing protein) containing proteotypic peptide DLMVQAGIENSIK (Pept-13). Illumina HiSeq and ONT MinION reads derived for HepG2 and liver (donor 1, liv.d1) samples, respectively, are mapped to the human genome (the visualization is created by Integrated Genome Viewer^53^). Homozygous substitution rs2298645 results in the disappearance of trypsin cleavage site and forming a longer peptide NVNDLMVQAGIENSIK (Pept-16).

The sequence of Pept-24 is split between exons 1 and 2 of the HSBP1L1 gene (Fig. 3), with the first two amino acids encoded at the tail of exon 1, while the rest of the peptide occupies the whole length of exon 2. The inspection of the splice region in Integrated Genome Viewer^53^ has shown no abnormalities that can course the variations in the sequence of this proteotypic peptide.

The second putative peptide of HSBP1L1 is located in exon 3. Fig. 3 provides an enlarged image of the corresponding genome region. The amino acid sequence encoded by exon 3 runs in a bottom line of the cartoon, with Pept-13 starting with “DLMV…” (the second-half part of the exon-spanning sequence). The two blocks laying over the sequence data correspond to the ONT and HiSeq reads obtained for the liver (donor 1) and HepG2 cells, respectively.

A noticeable finding depicted in Fig. 3 is G>T substitution at the position of the genome indicated by the vertical red rectangle. The observed G>T nonsynonymous SNP (rs2298645) has no clinical relevance and is characterized by ~81% of the overall population frequency as reported by various databases (1000 Genomes, ExAC, ALFA).

It is important to note that the substitution appears to be homozygous, and it causes the substitution of lysine (K) to asparagine (N) in the 55 position of HSBP1L1 sequence (K55N). In turn, the K55N substitution leads to the disappearance of the cleavage site, thus making the second proteotypic peptide of HSBP1L1 longer by 3 additional amino acid residues from the N-terminus (Pept-16).

We tried to search the MS/MS repositories for the aforementioned peptides – Pept-24 (*DAAENLFQELQEHFQALTATLNLR*), Pept-13 (*DLMVQAGIENSIK*) and Pept-16 (*NV[K>N]DLMVQAGIENSIK*, with K55N substitution). The MS/MS data were downloaded for different human cell lines ^32^. We also scouted our proprietary data^39^ including presently unpublished 2D-LC-Orbitrap dataset, as well as the results of proteomic profiling of A-549 cell line performed during COVID-19 study^54^. The library of human protein sequences (downloaded from SwissProt and contained 42,360 entries) was extended with an additional sequence of HSBP1L1 carrying K55N substitution.

### 6. Following the ONT-highlighter across proteomic resources and search engines

With a modified sequence database, the search was implemented using SearchGUI^30^ with a parent mass tolerance of 7 ppm and a fragment mass tolerance of 20 ppm. After processing in PeptideShaker, for X!tandem there were 61.5 thousand peptide hits corresponding to 6.8 thousand protein groups in HepG2 cell line (73.0 thousand peptides and 7.7 thousand protein groups in LNCaP, respectively) and slightly less protein groups - for OMSSA and MS-GF+ searches. Quite similar results were observed when using MaxQuant of the version 1.6.12.0 (as compared to version 1.5.3.31 used in the original work^32^).

We controlled our search parameters using the previously observed 24 a.a.r. proteotypic peptide Pept-24 and observed its identification in HepG2 and LNCaP cell lines (according to the Peptide Accession entry PAp05383439). In the HepG2 cell line, Pept-24 was reliably identified by all search engines under study (by MS/MS match). In the LNCaP cell line, Pept-24 was identified by MS-GF+ only, observing a clear pattern, similar to the spectra deposited in PeptideAtlas by Kulak *et al.*^32^, while OMSSA and X!Tandem could not detect Pept-24 in the LNCaP MS/MS spectra collection. The MaxQuant search engine identified this peptide in the LNCaP cell line only by merging several spectra (“matching between runs” option in MaxQuant)^32^.

The hits for Pept-24 (uniquely corresponding to the missing C9JCN9 protein) were detected by MaxQuant at an FDR of 1% for both peptides and proteins^32^ according to PeptideAtlas entry. The precursor intensity of Pept-24 in HepG2 was 2.00·10^6^ compared with a lower signal 0.98·10^6^ for LNCaP. Thus, by using Pept-24 as a control, we located the HepG2 and LNCaP spectra as a source of evidence for HSBP1L1. In our searches by MaxQuant Pept-24 was absent from most cell lines under study (including LNCaP) except for HepG2 and RKO (Table 2).

**Table 2.**
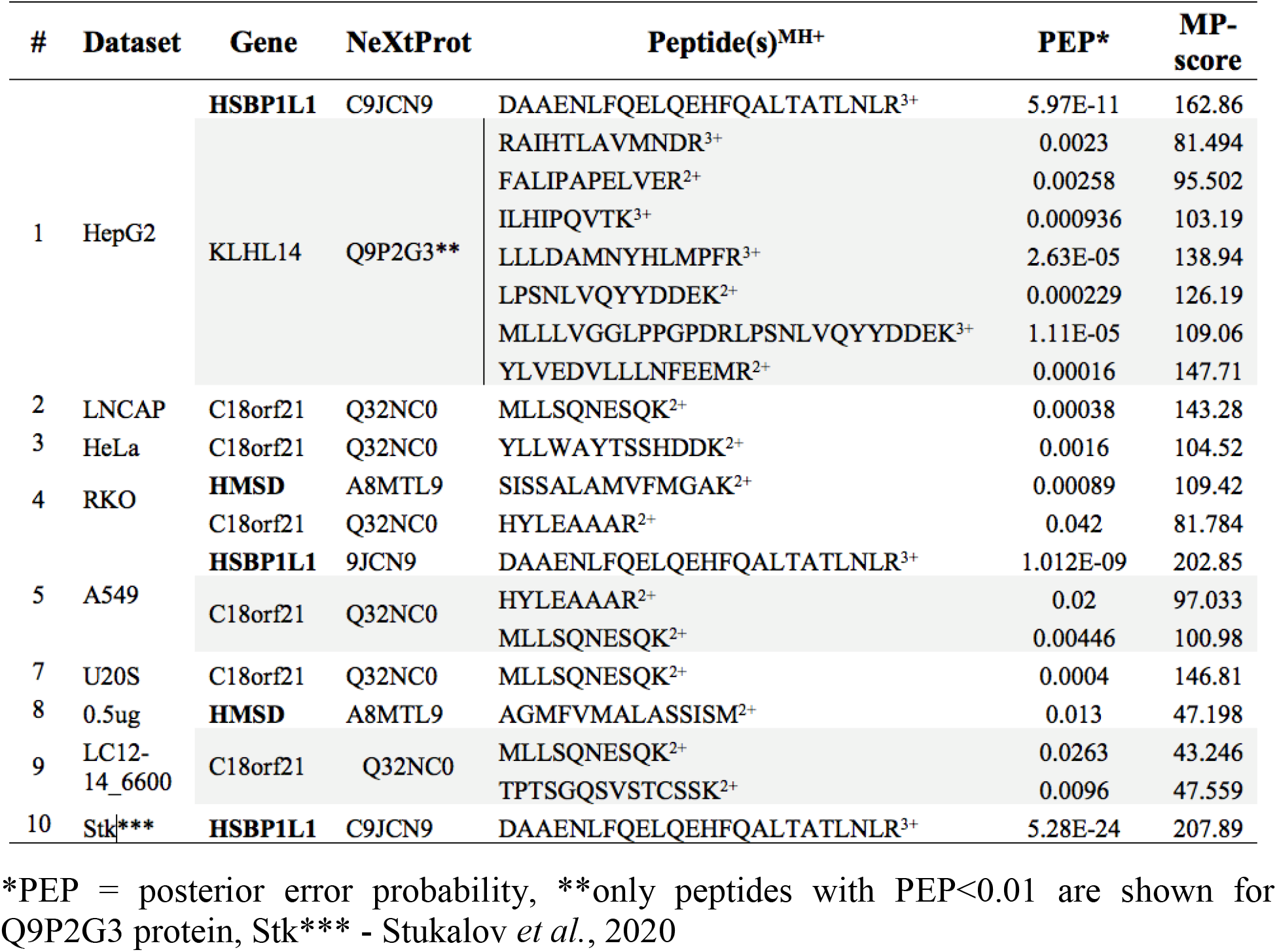
Occurrence of ONT-selected missing and uPE1 proteins across cell lines^32^, and SARS-CoV affinity purification^54^.

At the same time, the second peptide Pept-13 was missing from the datasets ^32^ for the cell lines and from our proprietary collection of MS/MS data^39^. In contrast to our expectations, the mutant version of the peptide (Pept-16) was also missing in these datasets.

Therefore, although HSBP1L1 was assigned to PE1 (“protein evidence at protein level”) since July 2020, we failed to reach the MS-based confirmation for this protein without using protein-specific enrichment methods, at least in the datasets of selected cell lines and our proprietary collection of MS/MS data.

Continuing the quest for MS-based evidence of the HSBP1L1 protein, the GPMdb resource^55^ was queried. Very weak evidence with mis-cleaved peptide 52-NV<u>K</u>DLMV[Q]AGIENSIK-68 (glutamine at the 60th position of the HSBP1L1 sequence is deamidated) was found with an expectation value of random match equal to 10^−2^ (for comparison, the e-value for the control Pept-24 was 10^−14^). Another finding was the 8 a.a.r.-long GPEAPGGR peptide, whose positions were at the beginning of the HSBP1L1 sequence (refer to the upper part of Fig. 3). The peptide matched to the product of the nonsense mediated decay of the HSBP1L1 gene, comprising 45 a.a.r. only (the K7ENV5 entry in UniProtKB). This peptide has the expectation values two orders of magnitude lower than abovementioned miscleavaged versions of Pept-13, but the length of the peptide still was one residue below the current threshold of the C-HPP Guidelines^43^.

In addition to the dataset provided by Kulak *et al.*, the GMBdb survey brought our attention to the dataset PXD020222, obtained by affinity purification mass spectrometry (Q-Exactive HF) analysis of SARS-CoV-2 and SARS-CoV proteins in A549 cells. HSBP1L1 (as well as C18orf21) was indicated in the protein groups report. For HSBP1L1, the already detected Pept-24 was observed with extremely high confidence, with an error probability (PEP) of 10^−75^ (the peptide score was 207). Two other HSBP1L1-specific peptides were identified with low confidence, with an error probability of 0.08 and scores of 20 and 83 for IEDLQKNVK and DLMVQAGIENSIK (Pept-13), respectively.

The uPE1 protein C18orf21 was reported by two peptides after affinity purification from A549 cells. The first peptide MLLSQNESQK produced a posterior error probability at an acceptable level of 0.00015 (score=124), and the second one was detected below the significance cutoff (PEP=0.03, score=70).

The SearchGUI package yielded no gain in determining the missing protein HSBP1L1 and other proteins listed in Table 1. That could be of misusage of the package by an author, wrong parameters etc., however it is quite illustrative why some proteins are still lost as a result of intuitively comfortable bioinformatics solutions.

Another peptide of HSBP1L1 was retrieved from the proteomicsdb.org as the MEEMGNRIEDLQK with Andromeda score 136 and a convincing expectation value of 10^−5^. This peptide is positioned in the protein sequence to the left-hand from the anchoring Pept-24.

With identyPROT search engine^37^ the two hits were observed in the APMS dataset^54^ obtained after fishing of proteins with the protein of coronavirus. The first hit Pept-24 occurred in the raw file labelled as COV2_APMS_5_4. The second peptide of 11 a.a.r. (GPEAPGGRALR) was located upstream to the Pept-24 and was identified in raw file, COV2_APMS_5_3, but not in COV2_APMS_5_4 (Fig. 4).

**Fig. 4.**
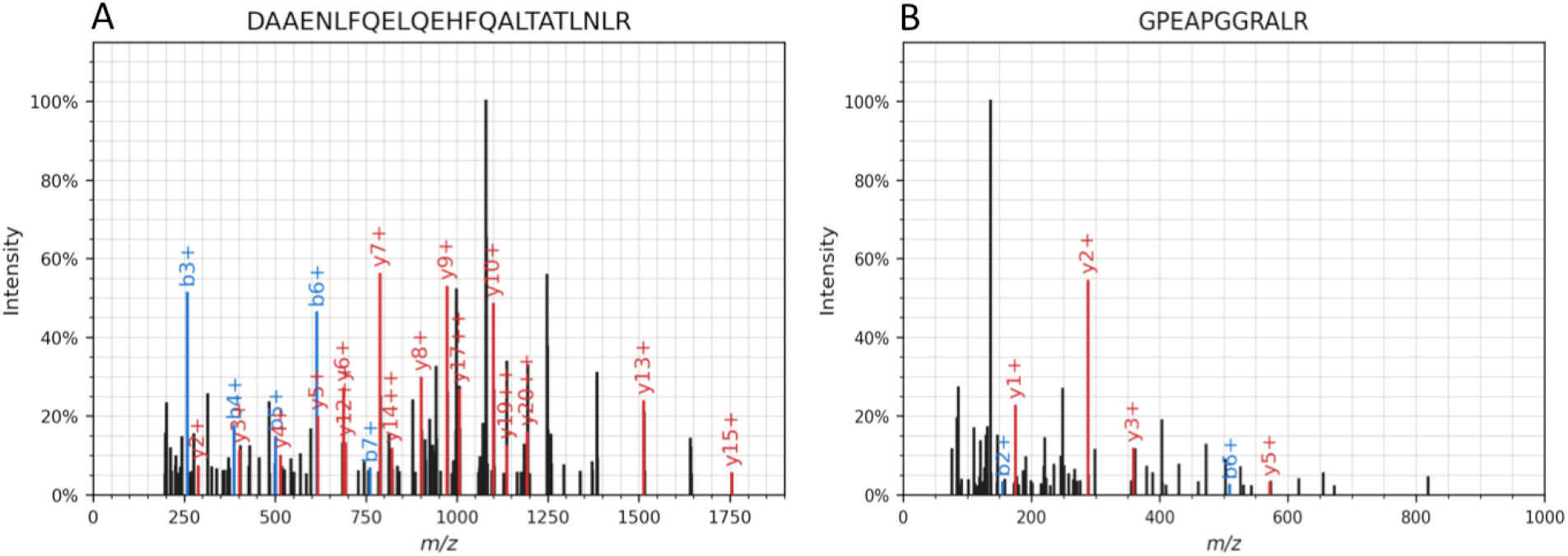
MS/MS of HSBP1L1 peptides identified by identyPROT search engine in the PXD020222 dataset^54^. Raw files: COV2_APMS_5_3 (A) and COV2_APMS_5_4 (B). The peptide GPEAPGGRALR did not passed the FDR < 0.01 cutoff. The same spectra (B) match with FDR <0.01 to the VYYEHLEK peptide from the NP_055318 protein (small subunit processome component 20 homolog) of 2785 aa.

The well-identified Pept-24 (charge 3+) scored e-value 10^−8^. For the second peptide the e-value was much weaker 10^−2^, failing to pass the FDR < 0.01 filter. There were no any signs of the Pept-13 and Pept-16, the former one derived from the transcriptome analysis (Fig. 3). In summary with other data from MaxQuant, the evidence of only the peptides of the first half of HSBP1L1 sequence were visible in several datasets and absent from the second half of the sequence, could indicate, that not the full-length protein, but the nonsense mediated decay product of HSBP1L1 (K7ENV5) was expressed, comprising only 45 a.a.r. In support of this thesis the confident identification in GPMdb referenced not only to the full-length sequence ENSP00000414236, but also referencing to the truncated version of the protein with identifier ENSP00000467108.

The GPEAPGGRALR peptide (spectra B at Fig. 4) could have problems in identification due to the peculiarities of its sequence. First the positive charge could be 1+. Secondly, the b-ions could be localized at the part of the spectra with low m/z values as the peptide is glycine-rich, thus producing the low-molecular weight fragment masses. Thirdly, the missed cleavage is present at the C-termini, while for instance the Pept-24 contains no missed-cleavages despite its severe length (spectra A at Fig. 4).

The information on selected set of proteins from dozen of cell lines pulled out by deep proteome survey^32^ or by the affinity purification^54^ was summarized in Table 2. The unique peptides with posterior error probabilities are shown. The C18orf21 being the uPE1 protein occurred more frequently, however mostly identified by a single peptide. The high abundance of KLHL14 in HepG2 as was shown by qPCR analysis (Table 1) was confirmed at the proteomic level with at least 4 peptides matching the current C-HPP guidelines.

Other target, the missing protein TMEM241, was absent from the datasets processed in this article (Table 2), In fact, in GPMdb/PRIDE also exist other datasets (e.g. PXD002815, PXD001333 and PXD014017), which were not used in this work, as experiments in Table 2 were quite illustrative that neither deep fractionation nor affinity purification can capture the second peptide for the missing HSBP1L1. The detection of this protein and other proteins no matter whether they were missing or uPE1 in most cases was limited by the presence of just a single peptide with the sufficiently high scoring, while the second peptides was observed as a rule as a statistically unjustified finding (Table 2). The signs of the target peptides were also lacking from our proprietary dataset PXD019263, acquired by applying 2D-LC-MS/MS to the HepG2 lysate [Vavilov et al., 2020, submitted]. Therefore, we can presumably assume for the Chr18 that the missing and uPE1 proteins share similarity because both types of proteins are successfully escaping the requirement of 2 peptides of 9 aa with FDR < 0.01 per protein (Table 2).

The data in Table 2 exemplifies the one-hit-wonders identifications of the HSBP1L1 and HSMD missing proteins. Surprisingly such problem was also shared by the studied uPE1s across cell lines (with an exception of KLHL14 confirmed by identification of 7 peptides in HepG2 cells only). It seems virtually impossible to gain the protein by MS-based technology in compliance with C-HPP Guidelines 3.0. At the same, cases listed in Table 2 provide the ground for the deeper inspection of the mass-spectrometry data enabling the chromosome-centric paradigm to limit analysis to just few valuable artifacts, likewise ascribed HSBP1L1 and C18orf21.

## Conclusions

The expression of approximately 50%–60% genes (depending on the biospecimen type) was detected simultaneously by qPCR, Illumina and Oxford Nanopore Technology (ONT)-based transcriptome sequencing (Supporting information Fig. S1). The most sensitive method of qPCR delivered the expression of 64% of the Chr18 genes in the liver tissue, and much more (77%) in HepG2. In total, 92% of the Chr18 coverage was achieved at the transcriptome level by compiling the data from different methods and only two types of biospecimens.

In the present work, we appended the ONT as the independent “third vote” to the previous pair of well-established methods such as qPCR and Illumina sequencing for transcriptomic research. We examined this in detail on an example of HSBP1L1 gene as a promising target for investigation. Our choice was further confirmed, because in the July 2020 version of NeXtProt HSBP1L1 has left the pool of missing proteins and gained the respectful PE1 status.

By analyzing jointly the missing and uPE1 proteins, we distinguished C18orf21 to study its function in the HepG2 cell line in the frame of CP50 challenge. The level of transcription of the gene is elevated up to 10 times in HepG2 cells compared with that in the liver (see Table 1). Another uPE1 protein – KLHL14 – demonstrated an even more striking mRNA expression profile, comprising 168 mRNA copies in a single HepG2 cell determined by RT–PCR, designating it a “Stakhanovite gene”, in contrast to the lack of expression in all three individual liver tissues.

We showed significant differences in the transcriptomic results obtained using different experimental methods: for many genes, they reached the order of magnitude. If we consider qPCR as the gold standard, RNA-Seq-based approaches may represent a reliable method only for qualitative evaluation of gene expression, whereas quantitative assessment may be significantly biased because of differences in sample preparation protocols and data processing pipelines. It was illustrated that, in connection with the problem of the detection of low-abundant transcripts, there was also the problem of approaching an actual picture of gene expression at the transcriptome level, unbiased by the sample preparation and data treatment procedures.

In this study, we showed the limitations of the correlation analysis of quantitative omics data, particularly, its strong dependency on the choice of correlation/clustering method. Spearman’s rank correlation analysis results are more susceptible to inaccuracies in the RNA-seq analysis of lowly expressed genes, but Pearson’s correlations are very prone to outliers for highly expressed genes. In any case, a great random component contributes to the results.

Finally, using the combination of short-read sequencing by Illumina HiSeq and long reads from ONT MinION, we have partially eliminated the suspicion that the problem of Chr18 missing proteins is associated with the choice of proteotypic peptides that are located in genomic regions containing nonsynonymous SNPs or nonannotated tissue-specific splice junctions. At the same time, we observed that, in the liver/HepG2 cells, the level of expression and read coverage of investigated missing proteins were comparable to those of other genes annotated as having protein evidence 1.

In conclusion, we illustrated the method that consists of obtaining the general chromosome-centric picture of the transcriptome datasets from different sources, followed by the analysis of highly expressed genes and transcripts for the functionally uncharacterized and missing proteins, with further verification of the proteotypic peptides of missing and uPE1 proteins by mapping onto the RNA-seq short and long reads.

## Supporting information

Table S1 - Quantitative data for the Liver and the HepG2 from several datasets

Table S2 - Stakhanovite genes of Chr18

Table S3 - Chr18-centric dendrogram-coupled correlation matrix

## Acknowledgments

The bioinformatics work was supported by the Russian Science Foundation grant #20-14-00328, using the equipment of the EIMB RAS “Genome” center (http://www.eimb.ru/rus/ckp/ccu_genome_c.php). The authors are grateful to the “Human Proteome” Core Facility, Institute of Biomedical Chemistry (IBMC) for performing PCR and RNA-Seq (ONT) and to Genotek Ltd. (Moscow, Russia) for performing RNA-Seq (Illumina).

## Supporting Information

**Contents**

**Supplementary Table S1.** – Quantitative data for the Liver and the HepG2 from several datasets, obtained in 2013 and 2020 with SOLiD, Illumina GII/HiSeq, qPCR and Oxford Nanopore (MinION) and mapped to the Chr18-encoded proteins

**Supplementary Table S2.** The top most expressed (“Stakhanovite”) genes of Chr18

**Supplementary Table S3.** – Chr18-centric dendrogram-coupled correlation matrix for different biosample types/years, sample preparation methods, transcriptome analytical platforms and bioinformatics pipelines

**Fig. S1.** Venn’s diagram for transcripts with TPM/FPKM values ≥ 0.1 obtained by qPCR and RNA-seq technologies.

**Fig. S2.1.** Correlations between transcripts’ abundances for 235 transcripts encoded on Chr18 and detected in both HepG2 cells and each of three liver samples (donors #1, #3, and #5) by qPCR analysis.

**Fig. S2.2.** Correlations between transcripts’ abundances for 121 transcripts encoded on Chr18 and detected in both HepG2 cells and each of three liver samples (donors #1, #3, and #5) by Illumina HiSeq sequencing.

**Fig. S3**. Splicing structure (Sashimi plots) of the observed at protein level but functionally uncharacterized (uPE1) gene C18orf21 derived by the Illumina/HiSeq (a) and Oxford Nanopore Technology, MinION (b) for the liver sample from donor #1.

## Notes

### Competing Interest Statement

The authors have declared no competing interest.

### Summary of Updates

In Supplementary file Figure S3 and S5 were removed and the appropriate corrections were made to the manuscript file.

