## Supplementary Figures for "Human Chr18: “Stakhanovite” Genes, Missing and uPE1 Proteins in Liver Tissue and HepG2 Cells"

**Contents**

**Supplementary Table S1.** – Quantitative data for the liver and the HepG2 cells from several datasets, obtained in 2013 and 2020 with SOLiD, Illumina GII/HiSeq, qPCR and Oxford Nanopore (MininION), and mapped to the Chr18-encoded proteins

**Supplementary Table S2.** The top most expressed ("Stakhanov") genes of Chr18.

**Supplementary Table S3.** Human Chr18-centric dendrogram-coupled correlation matrix for different biosample types/years, sample preparation methods, transcriptome analytical platforms and bioinformatics pipelines.

**Fig. S1.** Venn's diagram for transcripts with TMP/FPKM values ≥ 0.1 obtained by qPCR and RNA-seq technologies.

**Fig. S2.1.** Correlations between transcripts' abundances for 235 transcripts encoded on Chr18 and detected in both HepG2 cells and each of three liver samples (donors #1, #3, and #5) by qPCR analysis.

**Fig. S2.2.** Correlations between transcripts' abundances for 121 transcripts encoded on Chr18 and detected in both HepG2 cells and each of three liver samples (donors #1, #3, and #5) by Illumina HiSeq sequencing.

**Fig. S3**. Splicing structure (Sashimi plots) of the observed at protein level but functionally uncharacterized (uPE1) gene C18orf21 derived by the Illumina/HiSeq (a) and Oxford Nanopore Technology, MinION (b) for the liver sample from donor #1.

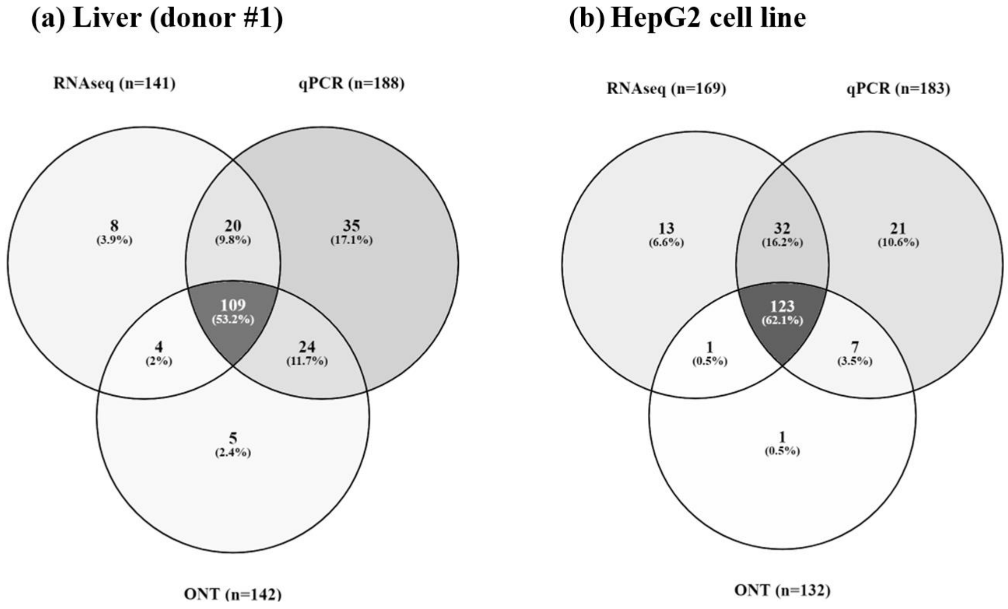


**Fig. S1.** Venn's diagram for transcripts with TMP/FPKM values ≥ 0.1 obtained by qPCR and RNA-seq technologies in **(a)** Liver tissue (number of detected transcripts = 205) and **(b)** HepG2 cell line (number of detected transcripts = 198). Visualized using Venny 2.1 (https://bioinfogp.cnb.csic.es/tools/venny/, Oliveros, J.C. (2007-2015) Venny: An interactive tool for comparing lists with Venn's diagrams.)

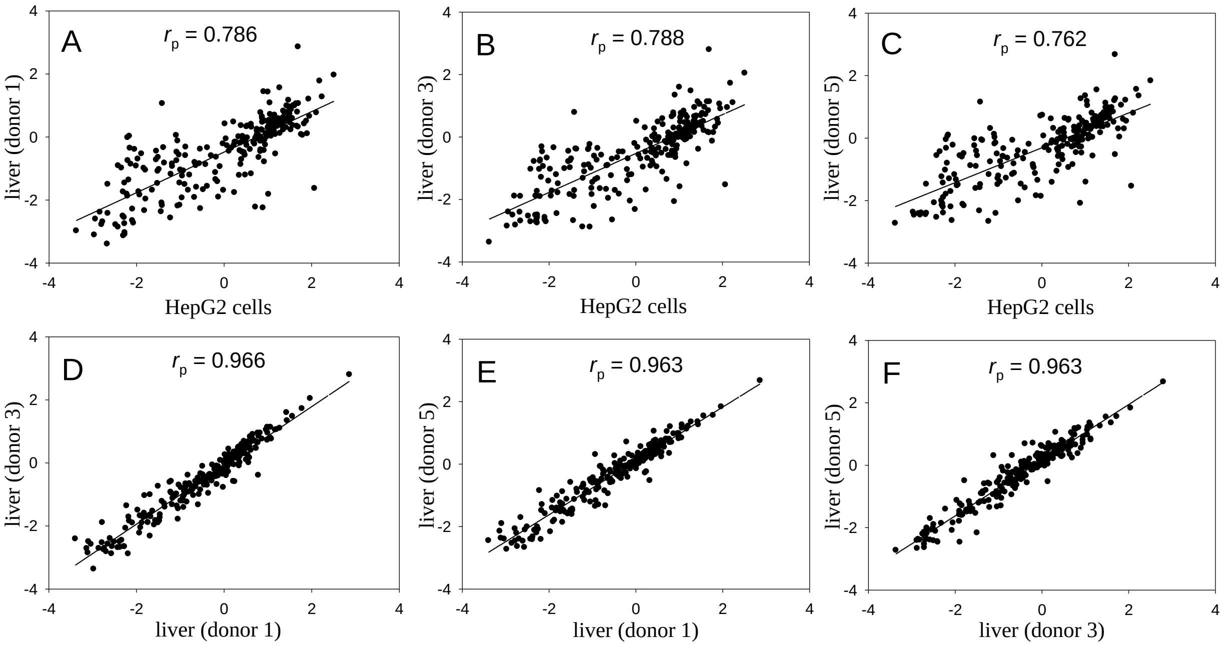


**Fig. S2.1.** Correlations between transcripts' abundances for 235 transcripts encoded on Chr18 and detected in both HepG2 cells and each of three liver samples (donors #1, #3, and #5) by qPCR analysis. Axis represent decimal logarithms of transcripts abundance. The transcript abundance is estimated as the copy numbers of cDNA per a cell. Values of Pearson's correlation coefficient, $r_p$, are shown in panels.

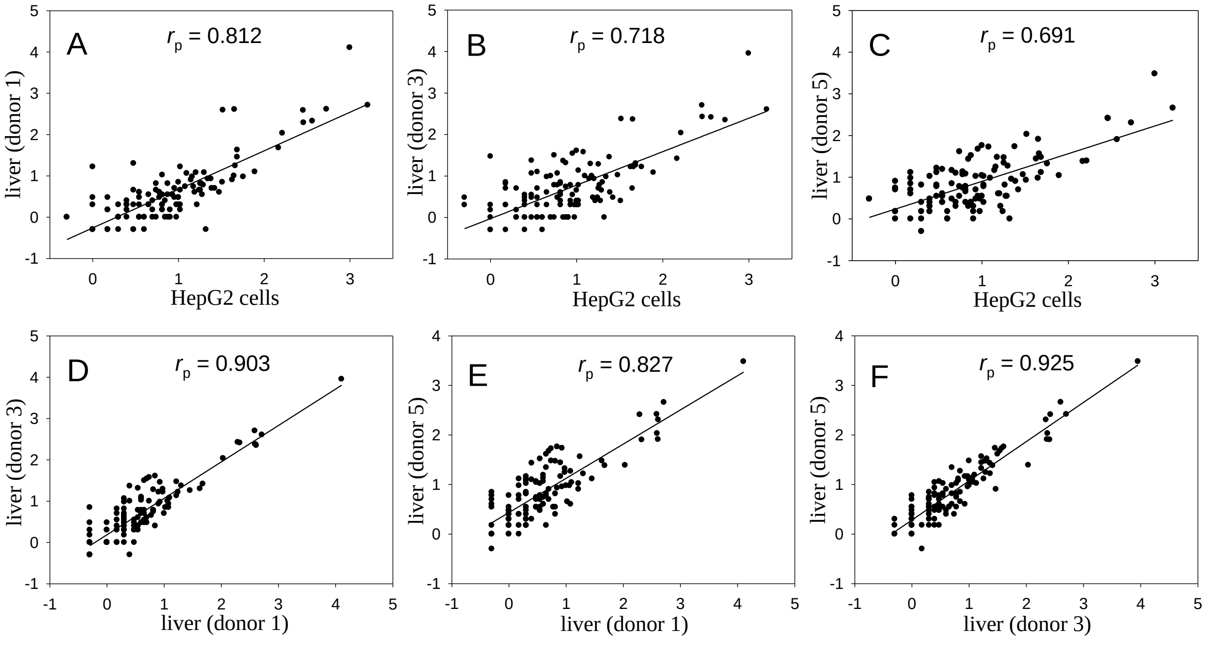


**Fig. S2.2.** Correlations between transcripts' abundances for 121 transcripts encoded on Chr18 and detected in both HepG2 cells and each of three liver samples (donors #1, #3, and #5) by Illumina HiSeq sequencing. Axes represent decimal logarithms of transcripts abundance. The transcript abundance is estimated as FPKM. Values of Pearson's correlation coefficient, $r_p$, are shown in panels.

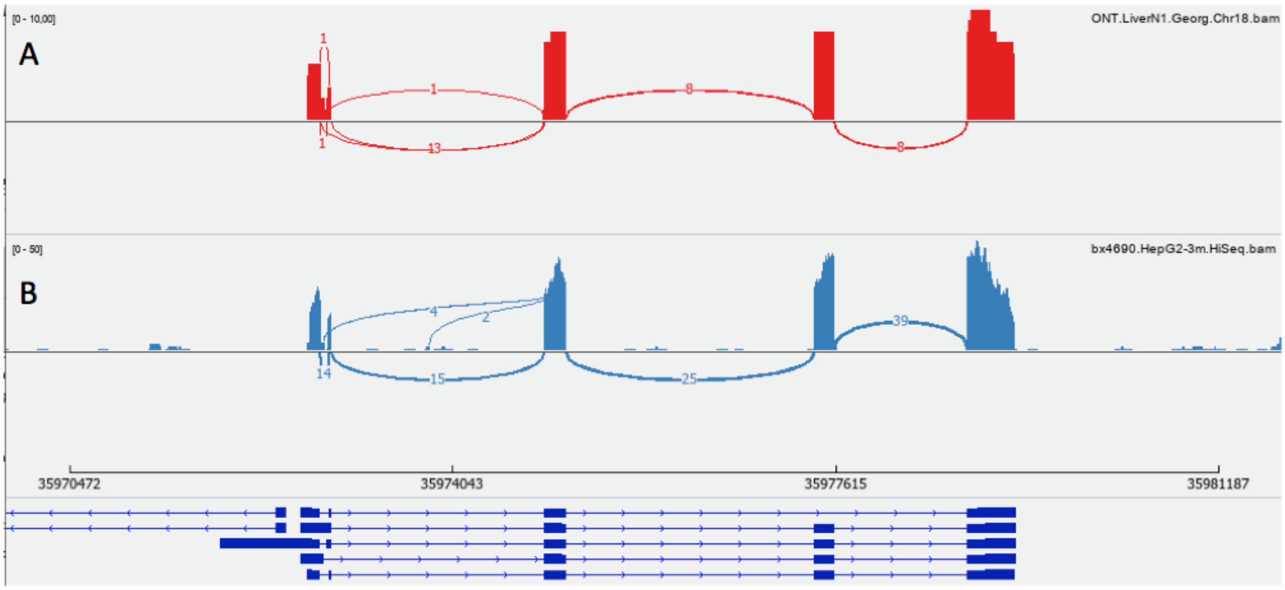


**Fig. S3**. Splicing sketch (Sashimi plot) of the observed as uPE1 (rarely seen at the protein level and functionally uncharacterized) translated from the C18orf21 gene. Derived by the Illumina/HiSeq (A) and Oxford Nanopore Technology, MinION (B) for the liver sample from donor #1.
